# Stable kinetochore–microtubule attachments restrict MTOC position and spindle elongation in acentrosomal oocytes

**DOI:** 10.1101/2020.05.07.081786

**Authors:** Aurélien Courtois, Shuhei Yoshida, Tomoya S. Kitajima

## Abstract

In mouse oocytes, acentriolar MTOCs functionally replace centrosomes and act as microtubule nucleation sites. Microtubules nucleated from MTOCs initially assemble into an unorganized ball-like structure, which then transforms into a bipolar spindle carrying MTOCs at its poles, a process called spindle bipolarization. In mouse oocytes, spindle bipolarization is promoted by kinetochores but the mechanism by which kinetochore–microtubule attachments contribute to spindle bipolarity remains unclear. This study demonstrates that the stability of kinetochore–microtubule attachment is essential for confining MTOC positions at the spindle poles and for limiting spindle elongation. MTOC sorting is gradual and continues even in the metaphase spindle. When stable kinetochore–microtubule attachments are disrupted, the spindle is unable to restrict MTOCs at its poles and fails to terminate its elongation. Stable kinetochore fibers are directly connected to MTOCs and to the spindle poles, and thus may serve as a measure that defines proper spindle length. These findings reinforce the hypothesis that kinetochores act as scaffolds for acentrosomal spindle bipolarity.

## Introduction

Spindle bipolarity is a prerequisite for chromosome segregation. In animal somatic cells, two centrosomes act as major microtubule nucleation sites and provide a spatial cue for bipolar spindle formation. In contrast, in acentrosomal cells, such as oocytes, a bipolar spindle forms without canonical centrosomes (Bennabi et al., 2016; Dumont and Desai, 2012; Howe and FitzHarris, 2013; Mogessie et al., 2018; Ohkura, 2015; Radford et al., 2017; Reber and Hyman, 2015). In mouse oocytes, acentriolar microtubule organizing centers (MTOCs) functionally replace centrosomes and serve as major microtubule nucleation sites (Blerkom, 1991; Clift and Schuh, 2015; Maro et al., 1985; Schuh and Ellenberg, 2007). The cytoplasm of oocytes initially carry many MTOCs, which are relocated around chromosomes upon nuclear envelope breakdown (NEBD). Microtubule nucleation from MTOCs leads to the assembly of an apolar microtubule-based structure called a microtubule ball, which then elongates into a bipolar-shaped spindle. Concomitant with the spindle elongation, MTOCs are sorted and relocated into the forming spindle poles (Clift and Schuh, 2015; Schuh and Ellenberg, 2007). These processes establish a bipolar-shaped spindle carrying MTOCs at its two poles.

Spindle elongation and MTOC sorting, two characteristic processes involved in spindle bipolarization, require the concerted action of microtubule regulators (Letort et al., 2019), including the antiparallel microtubule motor Kif11 (Mailhes et al., 2004; Schuh and Ellenberg, 2007), the microtubule bundle stabilizer HURP (Breuer et al., 2010), the spindle pole-focusing factor NuMA (Kolano et al., 2012), the minus end-directed motor HSET (Bennabi et al., 2018), and the Ndc80 complex (which recruits the antiparallel microtubule crosslinker Prc1 to kinetochores in an oocyte-specific manner) (Yoshida et al., in press). Despite increasing knowledge about the molecular mechanisms involved in establishing spindle bipolarity, less is known about how the bipolarity is maintained. Maintenance of spindle bipolarity is important to understand as its instability is a hallmark of error-prone human oocytes (Haverfield et al., 2017; Holubcová et al., 2015).

Kinetochore–microtubule attachment contributes to the integrity of spindle bipolarity. In centrosomal mitotic cells, the disruption of kinetochore–microtubule attachments can perturb spindle bipolarity, which is pronounced when centrosomal functions are impaired (Lončarek et al., 2007; Moutinho-Pereira et al., 2013; O’Connell et al., 2009; Toso et al., 2009). In mouse oocytes, mutations and knockdowns that perturb kinetochore–microtubule attachment are often associated with spindle bipolarity defects (Gui and Homer, 2013; Sun et al., 2010, 2011; Woods et al., 1999). However, the precise manipulation of the stability of kinetochore–microtubule attachments has not been tested. Unlike somatic cells, where sister kinetochore biorientation and entry to metaphase abruptly stabilize kinetochore-microtubule attachments, in oocytes, the stabilization of kinetochore–microtubule attachment is gradual throughout prometaphase and metaphase (Brunet et al., 1999; Davydenko et al., 2013; Kitajima et al., 2011; Yoshida et al., 2015). How the stabilization of kinetochore–microtubule attachment affects the dynamics of spindle elongation and MTOC sorting has not been quantitatively analyzed.

In this study, we show that the stability of kinetochore–microtubule attachments is required to confine MTOCs at spindle poles and to limit spindle elongation. Our quantitative analysis demonstrates that MTOC sorting is a gradual process. During metaphase, while the majority of MTOCs are positioned at the poles of the bipolar-shaped spindle, a small population of MTOCs are found in the central region of the spindle until they are eventually sorted to the poles. Reducing the stability of kinetochore–microtubule attachments with a phospho-mimetic form of Ndc80, a major microtubule-anchoring protein at kinetochores, causes defects in restricting MTOCs at the spindle poles and in terminating spindle elongation. Stable kinetochore fibers (K-fibers) are directly connected to MTOCs and to spindle poles, suggesting their roles in restricting MTOC position and in limiting spindle elongation.

## Results

### MTOC sorting is gradual and frequently leaves small MTOCs in the central region of the bipolar-shaped spindle

To investigate acentrosomal spindle assembly during meiosis I, we used high-resolution imaging of MTOCs in mouse oocytes. To visualize the dynamics of MTOCs, we tagged the MTOC marker Cep192 (Clift and Schuh, 2015) with mNeonGreen (mNG-Cep192). We introduced RNAs encoding mNG-Cep192 and the chromosome marker H2B-mCherry into mouse oocytes at the germinal vesicle (GV, prophase I-like) stage and induced meiotic maturation *in vitro.* The dynamics of MTOCs and chromosomes were recorded with a confocal microscope throughout meiosis I (Supplementary Movie 1). We reconstructed the datasets into 3D and manually determined the axis of the spindle at every time point. Side views of the spindle were used to visualize the distribution of MTOCs and chromosomes along the forming spindle axis (Figure 1A). Quantitative analysis showed that the MTOCs gradually moved towards the poles of the bipolarizing spindle until 6 hours after NEBD, which coincided with the gradual chromosome congression towards the spindle equator (Figures 1B–1D). These observations are consistent with previous reports (Breuer et al., 2010; Clift and Schuh, 2015). We found that even in bipolar-shaped spindles during metaphase (4–6 hours after NEBD) (Figure 1E and Supplementary Figure 1), a small population of MTOCs were frequently positioned in the middle region of the spindle (Figure 1A and 1B). These ‘central MTOCs’ were observed in 79% of oocytes at early metaphase (4 hours after NEBD, n=21) and in 47% of oocytes at mid-metaphase (6 hours, n=21). Immunostaining of the MTOC marker pericentrin confirmed the presence of central MTOCs during metaphase (Figure 1F). These observations indicate that MTOC sorting is gradual and continues even in the bipolar-shaped spindle during metaphase.

**Figure 1:**
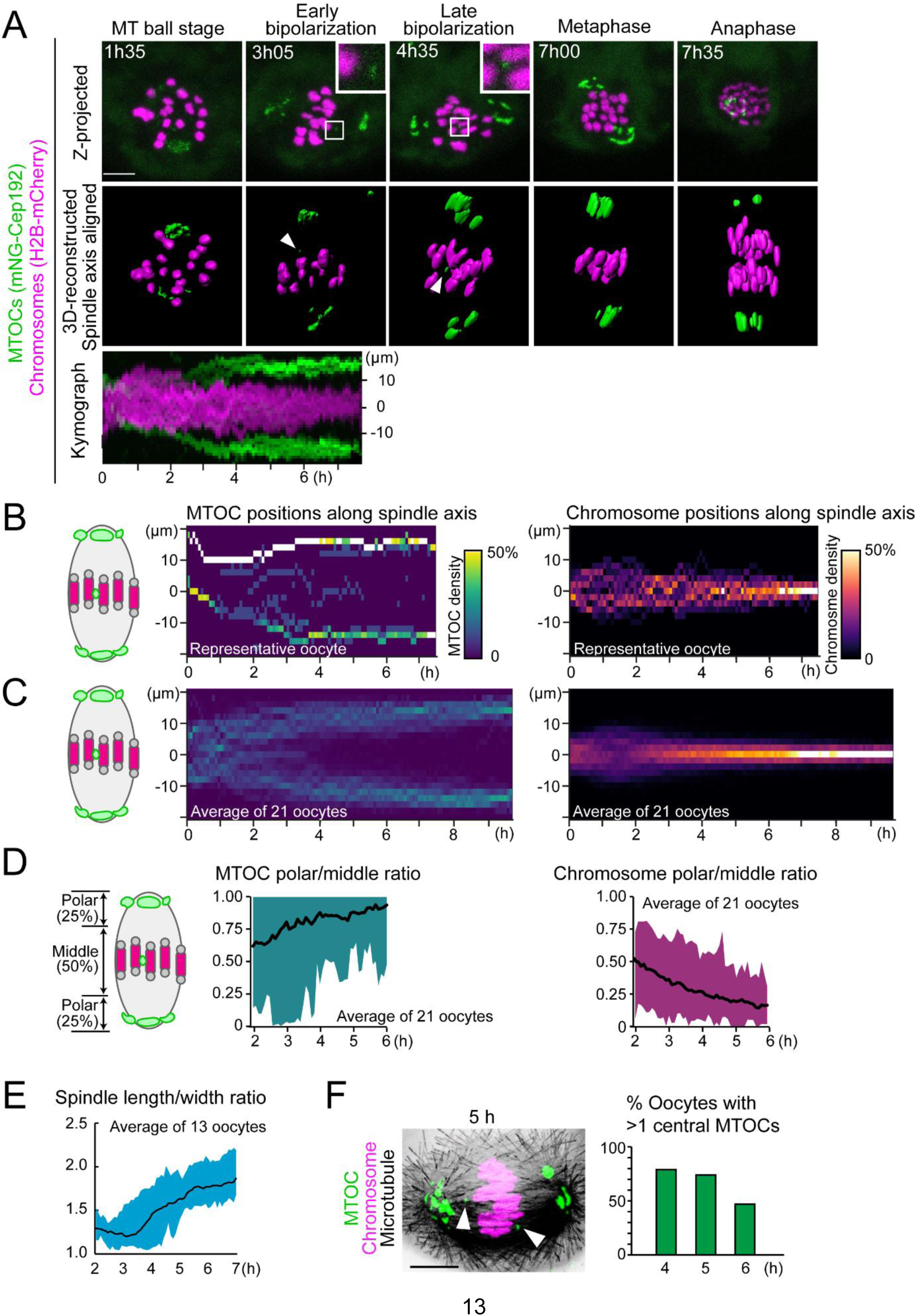
Establishment of 4D quantitative analysis for MTOC dynamics. **(A)** Live imaging of MTOC dynamics during meiosis I. Z-projection images (top) show MTOCs (mNG-Cep192, green) and chromosomes (H2B-mCherry, magenta). Central MTOCs are magnified (squares). 3D-reconstructed spindles are aligned with the spindle axis (middle). Central MTOCs are indicated with arrowheads. Time in h:mm. The kymograph (bottom) shows projected signals on the spindle axis for all time points. **(B)** Density maps of MTOCs (left) and chromosomes (right) along the spindle axis in a representative oocyte. The color represents the percentage of MTOCs or chromosome volumes, respectively, coded from dark (0%) to white (50% or more of the total volume). **(C)** Average density maps of 21 oocytes mapped as in (B). Data from 6 independent experiments are used. **(D)** Dynamics of MTOC sorting and chromosome congression. Ratio of MTOCs (left) or chromosomes (right) in the polar region versus those in the middle region of the spindle are plotted over time. **(E)** Dynamics of spindle elongation. The spindle was visualized with EGFP-Map4 (Supplementary Figure 1). The length/width ratio of the spindle was measured after 3D reconstruction (n=13 oocytes from 3 independent experiments). **(F)** Percentage of oocytes carrying one or more MTOCs in the middle region of the spindle. Oocytes were fixed and stained for MTOCs (pericentrin, green), chromosomes (Hoechst33342, magenta) and microtubules (α-tubulin, grey). n=21, 28, 24 from two independent experiments. A representative oocyte fixed at 5 hours after NEBD is shown. Arrowheads indicate central MTOCs. Time after NEBD. Mean ± SD are shown. Scale bars, 10 μm. See also Supplementary Movie 1.

### A majority of central MTOCs originate from polar regions of the forming spindle

To investigate the origin of central MTOCs, we recorded 4D datasets at a higher temporal resolution to track the dynamics of individual MTOCs (Figure 2A). We selected central MTOCs that were positioned in the middle of the spindle later than 4 hours after NEBD, and retrospectively analyzed their trajectories during prometaphase (Figure 2B). This analysis revealed that a majority of central MTOCs originated from the poles of the bipolar spindle (n=21/32, Figures 2C and 2D). These MTOCs spent substantial periods of time in the middle region of the spindle (an average of 69 minutes and a maximum of 210 minutes; n=21) (Figure 2C, Supplementary Movie 2), and then relocated either to the original (n=13/21) or to the opposite pole (n=8/21, Figures 2E and 2F). We found that a substantial number of the central MTOCs exhibited correlated motions with a closely positioned kinetochore (<2 μm) for considerable periods of time up to 60.5 minutes (Supplementary Figure 2, Movies 3 and 4), suggesting that central MTOCs can attach to kinetochores. These MTOCs eventually moved to the original pole or the opposite pole (Supplementary Figure 2, Movies 3 and 4), which suggests that central MTOCs can undergo a dynamic cycle of attachment and detachment to kinetochores. These observations indicate that MTOC sorting is highly asynchronous and includes active exchanges of MTOCs between the two forming spindle poles.

**Figure 2:**
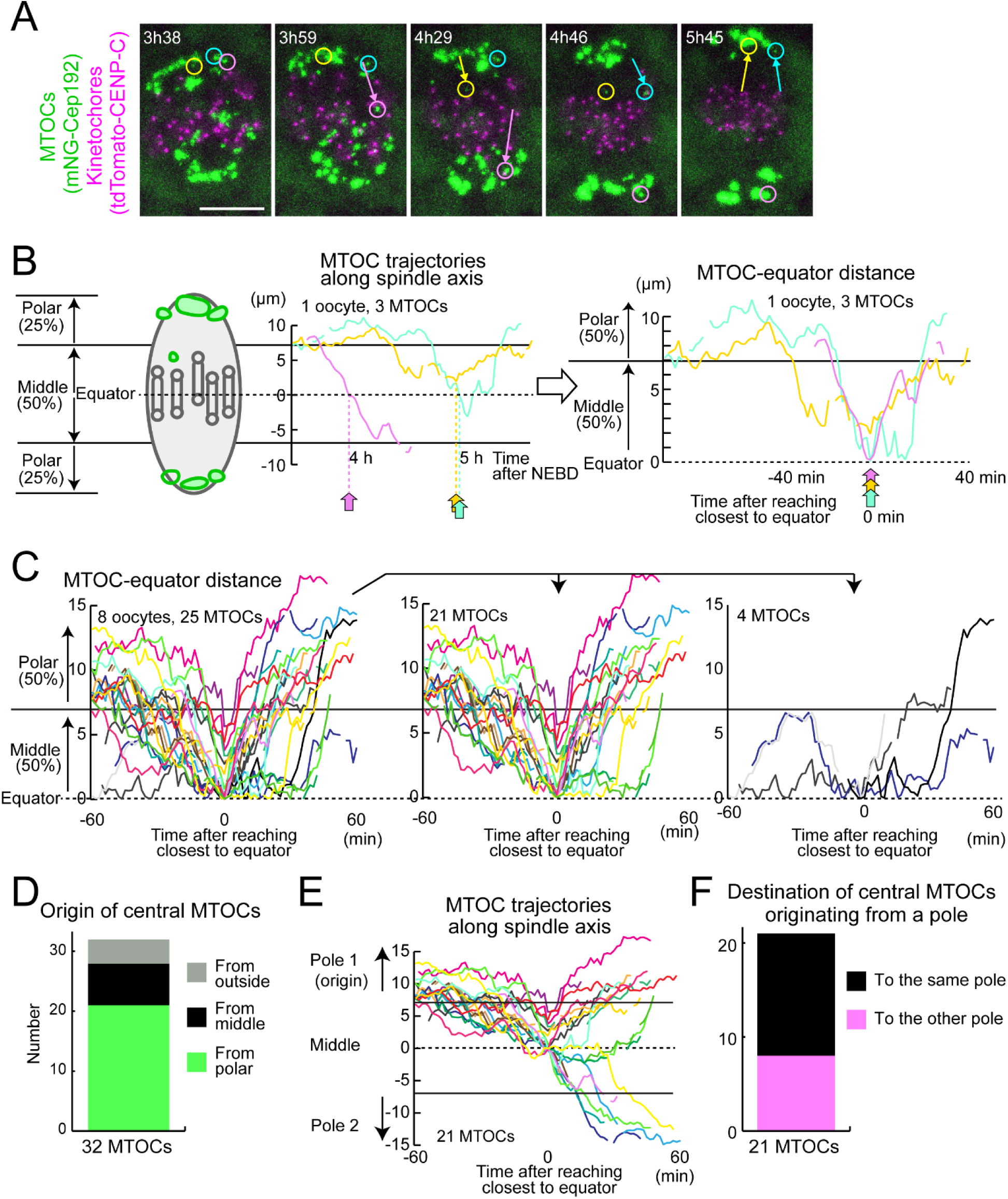
Central MTOCs originate from spindle poles. **(A)** Live imaging of MTOC and kinetochore dynamics. MTOCs (mNG-Cep192, green) and kinetochores (tdTomato-CENP-C, magenta) in BDF1 oocytes are shown. Time in h:mm after NEBD. Circles indicate central MTOCs, which are positioned in the middle region of the spindle later than 4 hours after NEBD. Arrows indicate displacement of the MTOCs. Scale bar, 10 μm. **(B)** Tracking analysis of central MTOCs. In the left graph, the positions of central MTOCs in (A) along the spindle axis are shown over time. MTOC positions were used to calculate distance to the spindle equator. In the right graph, lines showing temporal changes in MTOC-equator distance are aligned based on the time when the MTOC reached a position closest to the spindle equator. Horizontal lines at 7 and at −7 μm denote the thresholds used for the definition of the middle and polar regions of the spindle. **(C)** Central MTOCs originate from spindle poles and transiently stay in the middle region of the spindle. As in (B), the trajectories of central MTOCs (n=25 MTOCs of 8 oocytes from 3 independent experiments) were analyzed and their temporal changes in MTOC–equator distance are shown (Left graph). Central MTOCs were categorized into two groups: (1) ones that came from polar regions of the spindle (n=21, middle graph) and (2) others that stayed in the middle region throughout the period before reaching a position closest to the equator (n=4, right graph). **(D)** Origin of central MTOCs. Origins were categorized based on the results shown in (C). Note that 7 central MTOCs came from outside of the spindle and are not included in (C). **(E)** Destination of central MTOCs. The tracks of central MTOCs that originated from polar regions were used (n=21 MTOCs). The pole of origin was defined as Pole 1, while the other pole was defined as Pole 2. MTOC positions along the spindle axis (positive values for those closer to Pole 1) over time are shown. Note that central MTOCs moved back to the original pole (Pole 1) or switched their positions to the opposite pole (Pole 2). **(F)** Destination of central MTOCs that originated from a polar region was categorized as in (E). See also Supplementary Movie 2.

### Phospho-mutants of Ndc80 affect the stability of kinetochore-microtubule attachment

We recently reported that kinetochores promote acentrosomal spindle bipolarization during meiosis I (Yoshida et al., in press). We therefore hypothesized that kinetochore–microtubule attachments play a role in MTOC sorting. During meiosis I in oocytes, kinetochore–microtubule attachments become gradually stabilized by the dephosphorylation of kinetochore substrates including Ndc80, which is accompanied by the gradual conversion from relatively unstable lateral kinetochore–microtubule attachments to stable end-on attachments (Brunet et al., 1999; Davydenko et al., 2013; Kitajima et al., 2011; Yoshida et al., 2015). To manipulate the stability of kinetochore-microtubule attachments in oocytes, we replaced endogenous Ndc80 with the mutant versions Ndc80-9D (phospho-mimetic) and Ndc80-9A (phospho-deficient) (Guimaraes et al., 2008) by expressing those mutant forms in oocytes harvested from *Ndc80^f/f^ Zp3-Cre* mice (*Ndc80-* deleted oocytes) (Yoshida et al., in press) (Supplementary Figure 3A). To examine the stability of kinetochore-microtubule attachments, we visualized cold-stable microtubules at metaphase, a time when both lateral and end-on kinetochore-microtubule attachments were observed in control oocytes (Supplementary Figures 3B and 3C, “+ Ndc80-WT”). As expected, Ndc80-9D-expressing oocytes exhibited a significant decrease in end-on kinetochore–microtubule attachments (Supplementary Figures 3B and 3C, “+ Ndc80-9D”). Consistent with this observation, Ndc80-9D-expressing oocytes did not enter anaphase I by 14 hours after NEBD (n=25), while anaphase I entry was observed in 75% of Ndc80-WT-expressing oocytes by that time (n=33). In contrast, Ndc80-9A expression significantly increased end-on kinetochore-microtubule attachments and accelerated the onset of anaphase I (Supplementary Figure 3D). These results confirm that the stability of kinetochore-microtubule attachments is decreased in Ndc80-9D-expressing oocytes, whereas their stability is increased in Ndc80-9A-expressing oocytes.

### Phospho-mimetic Ndc80 is defective for restricting MTOC positions at spindle poles

Using Ndc80-9D and Ndc80-9A as tools to manipulate the stability of kinetochore–microtubule attachments, we addressed their effects on MTOC sorting during spindle bipolarization. Investigation of MTOC dynamics using our quantitative analysis pipeline revealed that Ndc80-9D-expressing oocytes failed to maintain MTOCs at the spindle poles (Figure 3A, Supplementary Movies 5 and 6). Although MTOCs tended to be positioned at the poles of the forming spindle during the early phases of MTOC sorting (2–4 hours after NEBD), they failed to maintain their positions at the spindle poles during the later phases (later than 4 hours) (Figures 3A and 3B). MTOCs that detached from the spindle poles frequently shuttled between the two poles (Figure 3B, Supplementary Movie 6), suggesting that spindle microtubules are dynamic but are incapable of properly sorting MTOCs. These defects resulted in the formation of spindles devoid of MTOCs at both poles (Figure 3C). In contrast to Ndc80-9D-expressing oocytes, Ndc80-9A-expressing oocytes did not exhibit any detectable differences from Ndc80-WT oocytes in MTOC sorting (Figures 3A–3C). These results indicate that Ndc80 dephosphorylation, which increases the stability of kinetochore–microtubule attachments, is required to restrict MTOC positions at spindle poles.

**Figure 3:**
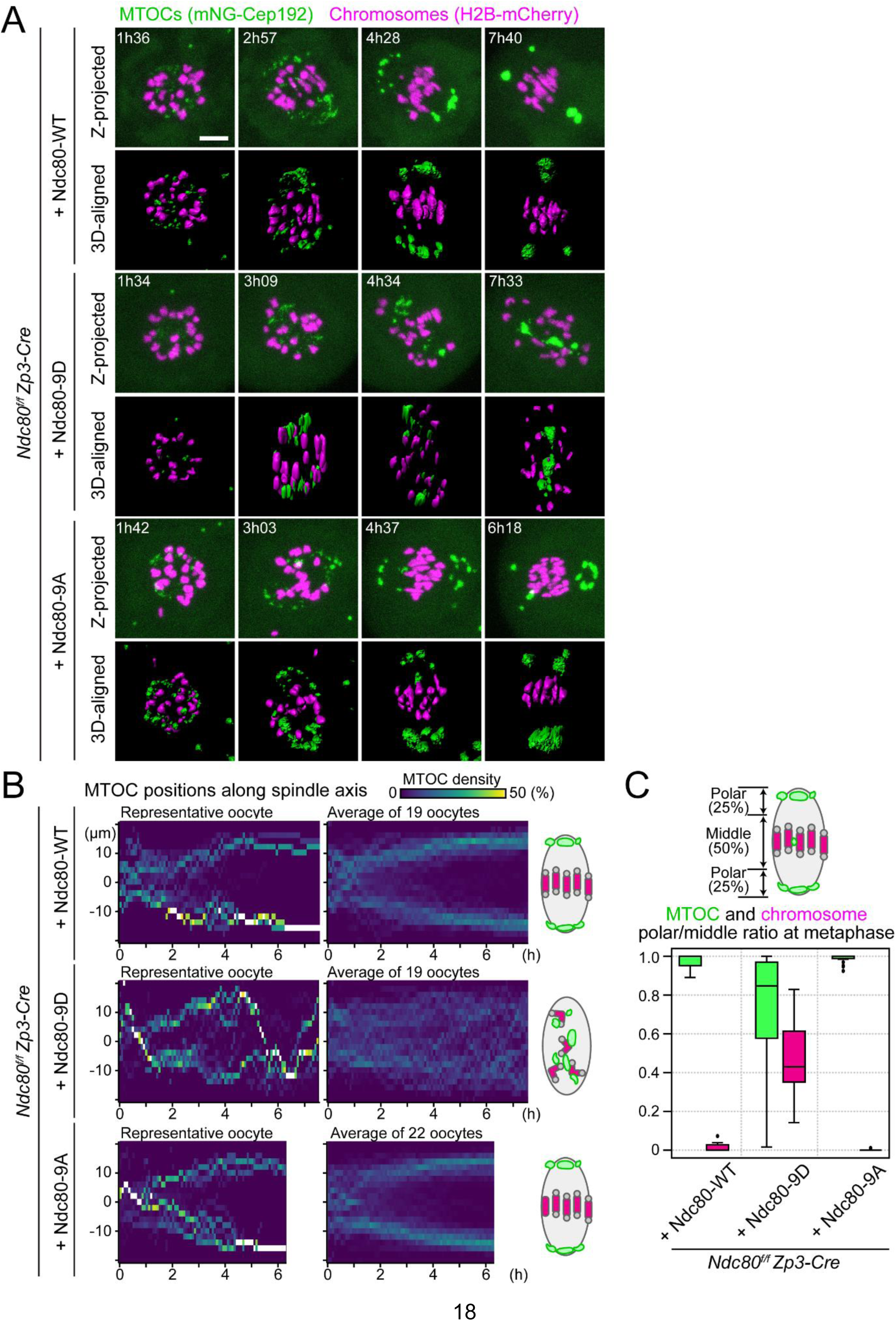
Kinetochore–microtubule attachment stability is required to confine MTOCs at spindle poles. **(A)** Live imaging of *Zp3-Cre Ndc80^f/f^* oocytes injected with Ndc80-WT, Ndc80-9D or Ndc80-9A. Z-projection and 3D-reconstructed images show MTOCs (mNeonGreen-Cep192, green) and chromosomes (H2B-mCherry, magenta). Time in h:mm after NEBD. Scale bar, 10 μm. **(B)** Ndc80-9D is defective for MTOC sorting. The density map of MTOCs along the spindle axis in *Zp3-Cre Ndc80^f/f^* oocytes expressing Ndc80-WT, Ndc80-9D or Ndc80-9A. The color represents the percentage of MTOC volume coded from dark (0%) to white (50% or more of the total volume). **(C)** MTOCs do not accumulate at the spindle poles in Ndc80-9D-expressing oocytes. Ratio of MTOCs (green) or chromosomes (magenta) in the polar region versus those in the middle region of the spindle at 6 hours after NEBD (n=19, 19 and 22 oocytes, respectively, from at least three independent experiments). See also Supplementary Movies 5 and 6.

### Dephosphorylation of Ndc80 is not required for initiating but is required for terminating spindle elongation

We next addressed whether the defects of Ndc80-9D for MTOC positioning are associated with spindle elongation defects, as centrosomes strongly affect spindle shape and bipolarity in centrosomal cells (Ganem et al., 2009). We quantified temporal changes in spindle shape in 3D with live imaging of the microtubule marker EGFP-Map4 (Schuh and Ellenberg, 2007) (Figure 4A, Supplementary Movie 7). Ndc80-9D-expressing oocytes had no significant delay in the onset of spindle elongation, but had significantly faster kinetics of elongation compared to Ndc80-WT- and Ndc80-9D-expressing oocytes (Figure 4B). Furthermore, in Ndc80-9D-expressing oocytes, the spindle failed to terminate its elongation after reaching a proper size (Figure 4B). The spindle length reached 37.3 ± 3.2 μm in Ndc80-9D-expressing oocytes at 7 hours after NEBD, which was significantly larger than spindle lengths observed in Ndc80-WT- and Ndc80-9A-expressing oocytes (32.3 ± 3.1 μm and 33.2 ± 3.0 μm, respectively) (Figure 4C). These results demonstrate that Ndc80 dephosphorylation is dispensable for making a bipolar-shaped spindle but is required for limiting spindle elongation.

**Figure 4:**
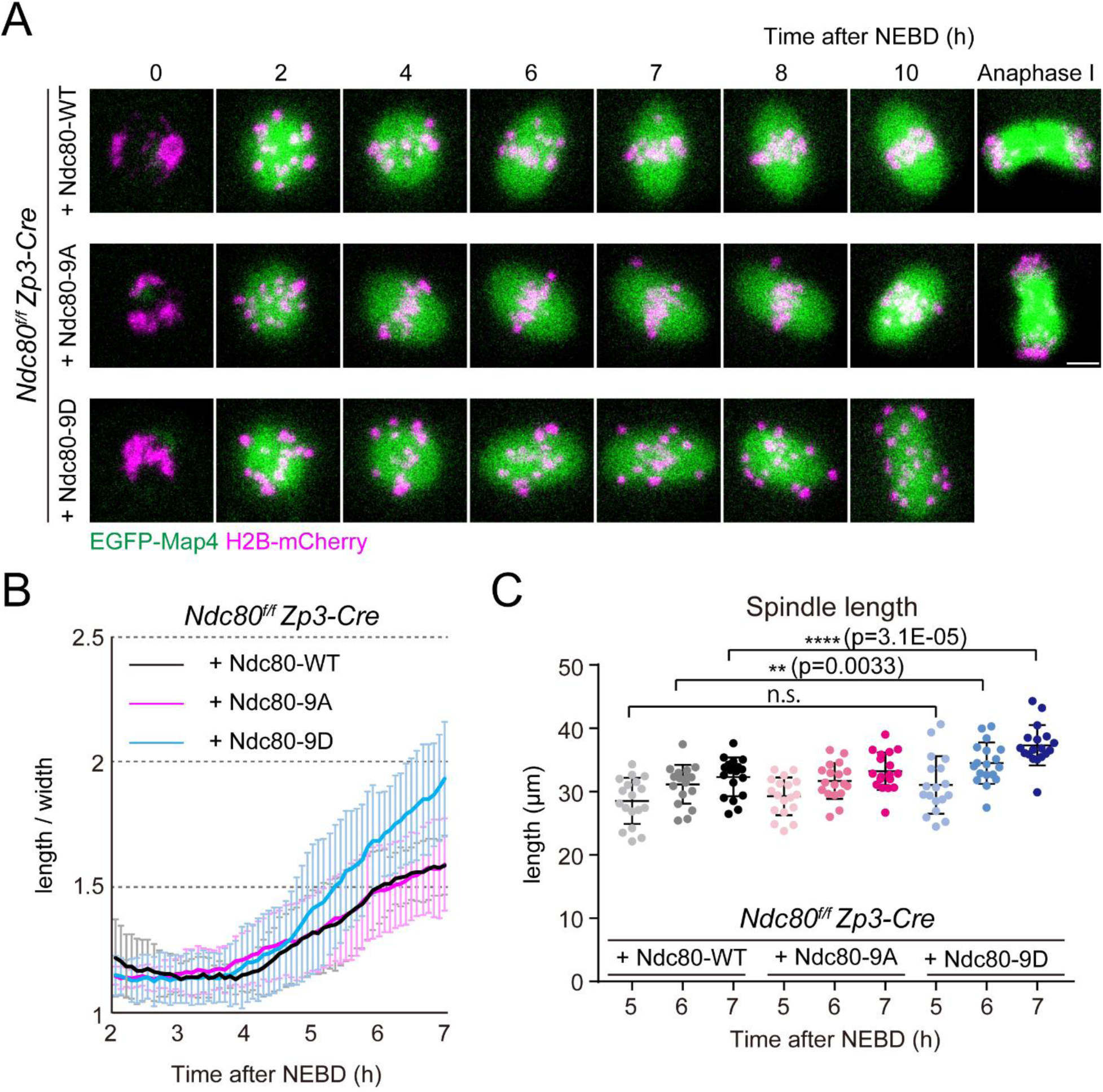
Kinetochore–microtubule attachment stability is not required for initiating but is required for terminating spindle elongation. **(A)** Live imaging of *Zp3-Cre Ndc80^f/f^* oocytes injected with Ndc80-WT, −9D or −9A. Z-projection and 3D-reconstructed images show microtubules (EGFP-Map4, green) and chromosomes (H2B-mCherry, magenta). Scale bar, 10 μm. **(B)** Ndc80-9D is defective for limiting spindle elongation. The aspect ratio (length/width) of 3D-reconstructed spindles was measured over time (n=18, 18 and 18 oocytes, respectively, from 3 independent experiments). **(C)** Ndc80-9D causes excessive spindle elongation. Spindle lengths measured after 3D reconstruction are shown. Scale bar, 10 μm. Means ± SD are shown. See also Supplementary Movie 7.

### The spindle in Ndc80-9D-expressing oocytes has a bipolar microtubule organization, despite the lack of MTOCs at the poles

Next, we tested whether bipolar-shaped spindles devoid of MTOCs at their poles, which are observed in Ndc80-9D-expressing oocytes, are capable of maintaining a bipolar microtubule organization. To quantify microtubule organization, we tagged the microtubule plus-end marker EB3 (Schuh and Ellenberg, 2007) with three copies of mEGFP (EB3-3mEGFP) and co-expressed it with the chromosome marker H2B-mCherry. The dynamics of microtubule plus-ends and chromosomes were recorded on a single-plane with confocal microscopy at 5 hours after NEBD (Supplementary Figure 4A and Movie 8). Our image processing pipeline allowed us to track microtubule plus-end comets, which was used to determine their directionality (toward one pole or the other) and position (the middle region or polar regions) in the spindle (Supplementary Figure 4A). That analysis revealed that the spindles of Ndc80-9D-expressing oocytes had a bipolar microtubule organization (Supplementary Figure 4B). In Ndc80-9D-expressing oocytes, the polar regions of the spindles exhibited significantly more comets moving toward the inside of the spindle (‘inward’), compared to those moving toward the spindle pole (‘outward’), as observed in control and in Ndc80-WT-expressing oocytes (Supplementary Figure 4B). Thus, the dephosphorylation of Ndc80 is not absolutely essential for the maintenance of bipolar microtubule organization in the spindle.

### Stable kinetochore fibers are directly connected to MTOCs and extend to the spindle poles

To gain insights into how stable kinetochore–microtubule attachments restrict MTOC position and spindle elongation, we investigated stable microtubules attached to kinetochores (K-fibers). High-resolution confocal imaging of metaphase spindles revealed that a substantial population of K-fibers were directly connected to MTOCs and extended to the end of the spindle poles (Figure 5A). Quantification demonstrated that 84 ± 9% of MTOCs were attached to K-fibers (Figure 5B). The average length of K-fibers was 11.2 ± 2.4 μm, which corresponded to 85 ± 17% of the distance between kinetochores and a spindle pole (Figure 5C). These results suggest that stable kinetochore–microtubule attachments regulate MTOC position and spindle elongation through K-fibers.

**Figure 5:**
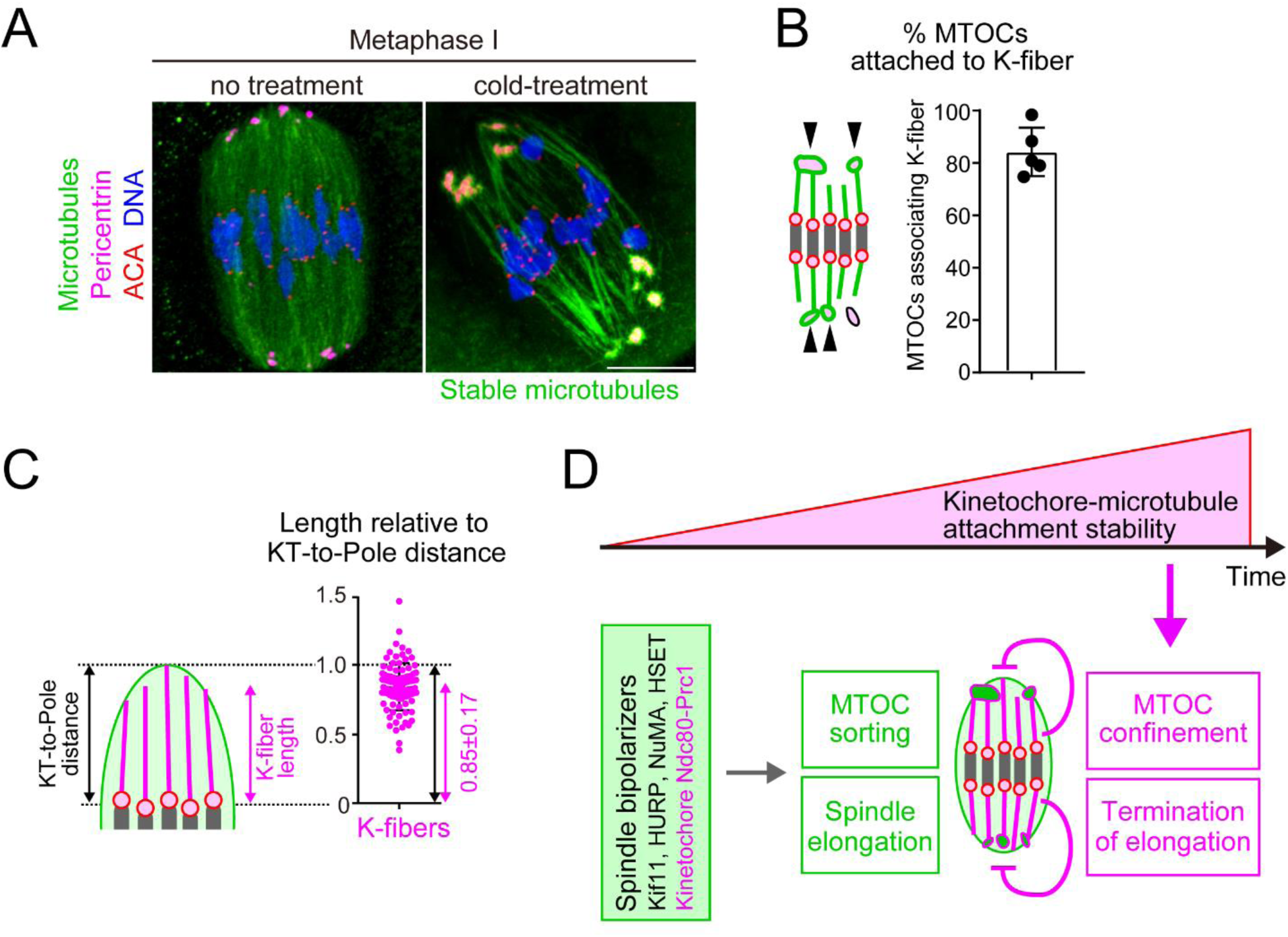
Model showing how kinetochore–microtubule attachments contribute to acentrosomal spindle bipolarization. **(A)** K-fibers are connected to MTOCs and extend to spindle poles. Oocytes 6 hours after NEBD were fixed following cold treatment. Microtubules (α-tubulin, green), MTOCs (pericentrin, magenta), kinetochores (ACA, red) and DNA (Hoechst333342) are shown. Five oocytes from 2 independent experiments were analyzed. Scale bar, 10 μm. **(B)** A majority of MTOCs are attached to K-fibers. The volumes of individual MTOCs were measured after 3D reconstruction. The sum volume of MTOCs attached to K-fibers relative to the total volume of MTOCs was calculated for each oocyte (n=5 oocytes from 2 independent experiments). **(C)** K-fibers extend to spindle poles. The length of K-fibers and distance between kinetochores and a spindle pole (KT-to-Pole distance) were measured. K-fiber length relative to the KT-to-Pole distance was calculated (n=90 K-fibers of 5 oocytes from 2 independent experiments). **(D)** Model summarizing the results of this study. In early phases of acentrosomal spindle assembly, spindle bipolarizers, including Kif11, HURP, NuMA, HSET, and the kinetochore Ndc80 complex that recruits Prc1, establish spindle bipolarity by promoting MTOC sorting and spindle elongation. Meanwhile, the stability of kinetochore–microtubule attachments gradually increases, depending on the dephosphorylation of kinetochore components including Ndc80. As spindle bipolarity is established, stable kinetochore–microtubule attachments spatially confine MTOCs at spindle poles and limit spindle elongation. Means and SD are shown.

## Discussion

The findings of this study, together with recent studies, suggest a model for how kinetochore– microtubule attachments contribute to acentrosomal spindle assembly (Figure 5D), which is triggered by the activation of microtubule nucleation from MTOCs. Nucleated microtubules initially form an apolar ball-like structure, which then undergoes spindle elongation depending on the action of microtubule regulators including Kif11, HURP, NuMA, HSET and the kinetochore Ndc80 complex that recruits Prc1 (Breuer et al., 2010; Kolano et al., 2012; Mailhes et al., 2004; Schuh and Ellenberg, 2007; Yoshida et al., in press). Those spindle bipolarizers promote MTOC sorting and spindle elongation. In parallel, the stabilization of kinetochore–microtubule attachments depends on dephosphorylation of the Ndc80 complex, which is promoted by Cdk1 activity that promotes the BubR1-mediated recruitment of PP2A-B56 phosphatase to centromeres (Davydenko et al., 2013; Yoshida et al., 2015). The stabilization of kinetochore–microtubule attachments is gradual, as Cdk1 activity gradually increases throughout prometaphase and metaphase (Brunet et al., 1999; Davydenko et al., 2013; Kitajima et al., 2011; Yoshida et al., 2015). Stable kinetochore–microtubule attachments start to appear when the spindle becomes bipolarized and thus initiates elongation (Davydenko et al., 2013; Kitajima et al., 2011; Yoshida et al., 2015). Spindle bipolarization allows the full stabilization of kinetochore–microtubule attachments by generating tension across the chromosome (Vallot et al., 2018; Yoshida et al., 2015). At later stages of spindle bipolarization, stable kinetochore–microtubule attachments spatially confine MTOCs at the poles of the spindle and terminate spindle elongation. Understanding how stable kinetochore–microtubule attachments mediate these functions requires future studies. K-fibers are distinctively long (36.5 ± 7.4 % of the spindle length) compared to typical spindle microtubules (5–27% of the spindle length in kinetochore-free extracts of *Xenopus* eggs) (Brugués et al., 2012), are directly connected to polar MTOCs and extend to the spindle poles. Stable kinetochore–microtubule attachments may enable K-fibers to anchor MTOCs, which may serve as a measure that properly adjusts the spindle length.

Bipolar-shaped spindles devoid of MTOCs at their spindle poles can maintain their overall bipolarity in shape and their microtubule organization. The dispensability of polar MTOCs for spindle bipolarity is in contrast to the essentiality of bipolar positioning of centrosomes for the spindle in centrosomal cells (Ganem et al., 2009). The dispensability of bipolar positioning of MTOCs for the bipolar shape of spindles is consistent with a previous report that the simultaneous inhibition of Kif11 and dynein allows the formation of bipolar-shaped spindles without MTOC splitting into two poles (Clift and Schuh, 2015). Although MTOCs are observed in oocytes of diverse animal models, including flies (Sköld et al., 2005), worms (Connolly et al., 2015), frogs (Gard, 1992), and mice (Blerkom, 1991; Maro et al., 1985; Schuh and Ellenberg, 2007), MTOCs have not been detected in human oocytes (Holubcová et al., 2015). In mouse oocytes, although microtubules are predominantly nucleated at MTOCs during the early stages of spindle assembly (prior to the microtubule ball stage) (Schuh and Ellenberg, 2007), microtubule nucleation in later stages may not be determined by MTOCs but are largely attributed to nucleation within the spindle, as observed by microtubule plus-end tracking in bipolar-shaped spindles (Supplementary Figure 4). Previous studies showed that defects in MTOC sorting are associated with spindle instability and chromosome segregation errors (Breuer et al., 2010; Kolano et al., 2012). It is thus likely that MTOC positioning at the spindle poles is critical for the robustness of acentrosomal spindles.

In a previous study, the expression of Ndc80-9A following morpholino-mediated knockdown of endogenous Ndc80 led to severe defects in spindle morphology (Gui and Homer, 2013). Such severe defects were not observed in our experimental system using Ndc80-9A expression in oocytes after Zp3-Cre-mediated *Ndc80* gene knockout. While that previous study depleted Ndc80 protein by culturing oocytes *in vitro* for 2 days after morpholino introduction, our study achieved protein depletion by maintaining oocytes *in vivo* for >10 days after gene knockout by Zp3-Cre. These apparent discrepancies may result from differences in depletion levels of Ndc80 protein or in the use of long-term *in vitro* cultures between the two experimental systems. Our observation that Ndc80 replacement with Ndc80-9D rather than with Ndc80-9A had detrimental defects in chromosome alignment is consistent with previous observations in somatic cells (DeLuca et al., 2011; Tooley et al., 2011).

MTOC sorting is gradual and lagging MTOCs are frequently observed in the central region of bipolar-shaped spindles during metaphase. The lagging MTOCs can be positioned close to kinetochores on bioriented chromosomes, underscoring the importance of active error corrections of kinetochore–microtubule attachments during metaphase (Lane and Jones, 2014; Yoshida et al., 2015). Lagging MTOCs could result from, and may facilitate, kinetochore–microtubule attachment errors and spindle instability, which are hallmarks of oocytes that lead to chromosome segregation errors (Holubcova et al., 2015). Lagging MTOCs thus may represent the error-prone nature of acentrosomal spindle assembly in oocytes.

We previously showed that the Ndc80 complex recruits the antiparallel microtubule crosslinker Prc1, which promotes spindle bipolarization, independently of kinetochore–microtubule attachments (Yoshida et al., 2020). Thus, the Ndc80 complex plays at least two roles in spindle bipolarity: (1) it promotes spindle bipolarization by recruiting Prc1, and (2) it confines MTOC positions at the spindle poles and limits spindle elongation by anchoring stable kinetochore– microtubule attachments. The findings of this study thus reinforce the hypothesis that kinetochores act as scaffolds for acentrosomal spindle bipolarity.

---

**Supplementary Figure 1:**
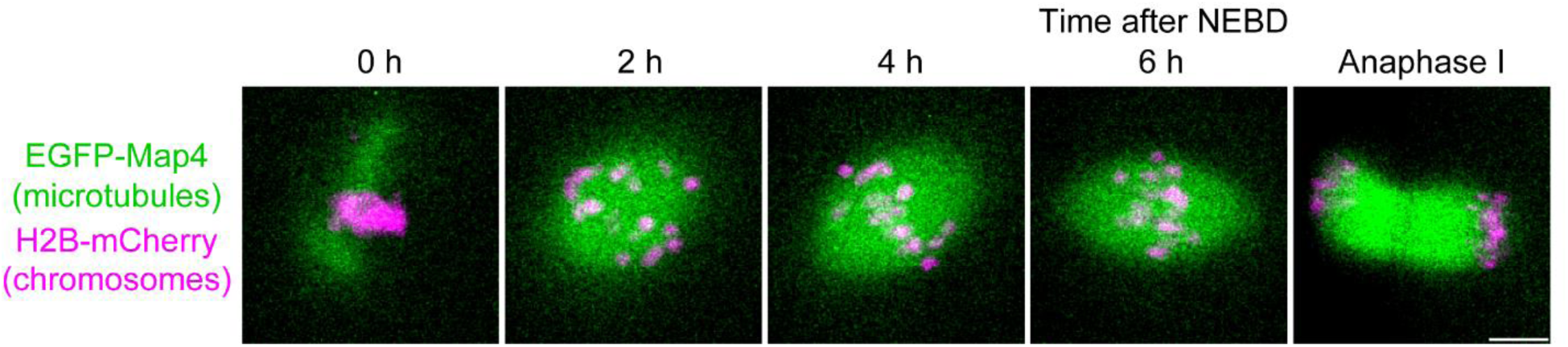
The kinetics of spindle elongation. Live imaging of oocytes expressing EGFP-Map4 (microtubules, green) and H2B-mCherry (chromosomes, magenta). Scale bar, 10 μm.

**Supplementary Figure 2:**
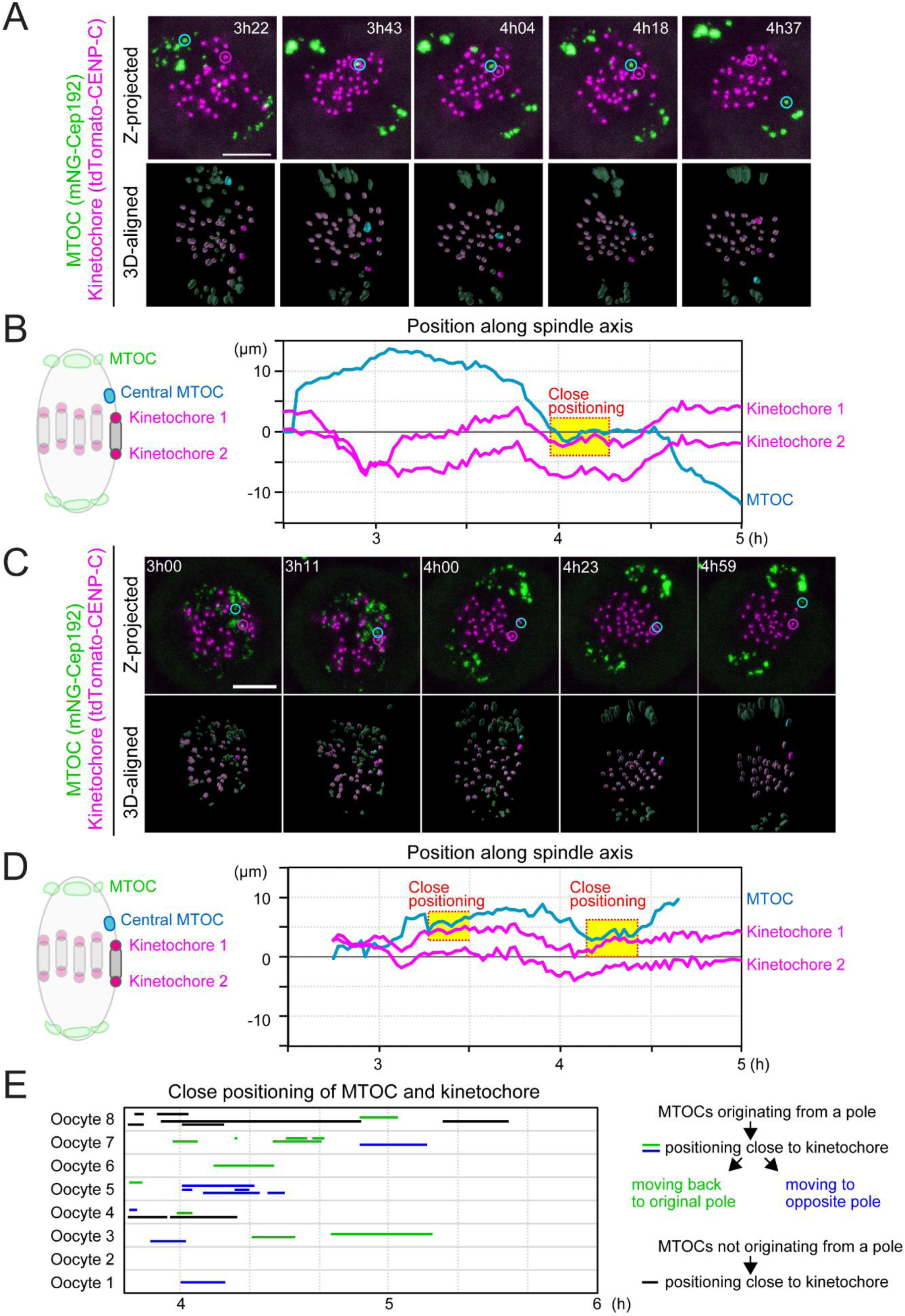
Central MTOCs can be positioned close to kinetochores. **(A and C)** Live-imaging of MTOC and kinetochore dynamics. Z-projected and 3D-reconstructed images for MTOCs (mNG-Cep192, green) and kinetochores (tdTomato-CENP-C, magenta) are shown. The 3D-reconstructed images are aligned based on the spindle axis, with highlights on an MTOC and a pair of kinetochores. Time in h:mm after NEBD. Scale bar, 10 μm. Cyan circles follow an MTOC in the middle of the spindle. Magenta circles follow a kinetochore that transiently positions close to the MTOC. **(B and D)** Tracking of MTOCs and kinetochores. The positions of MTOCs (cyan) and kinetochores (magenta), which are highlighted in (A) and (C), along the spindle axis are shown over time. Time after NEBD. Dotted boxes represent periods when the kinetochore-MTOC distance was less than 2 μm. **(E)** Close positioning of MTOCs and kinetochores is frequently observed. Lines indicate periods when oocytes exhibited close positioning of MTOCs and kinetochores (<2 μm). We categorized kinetochore-proximal MTOCs into three groups: (1) MTOCs that originated from a pole, positioned close to a kinetochore, and then moved back to the original spindle pole (green), (2) MTOCs that originated from a pole, positioned close to a kinetochore, and then moved to the opposite spindle pole (blue). (3) MTOCs that did not originate from spindle poles and are positioned close to the kinetochore (black). See also Supplementary Movies 3 and 4.

**Supplementary Figure 3:**
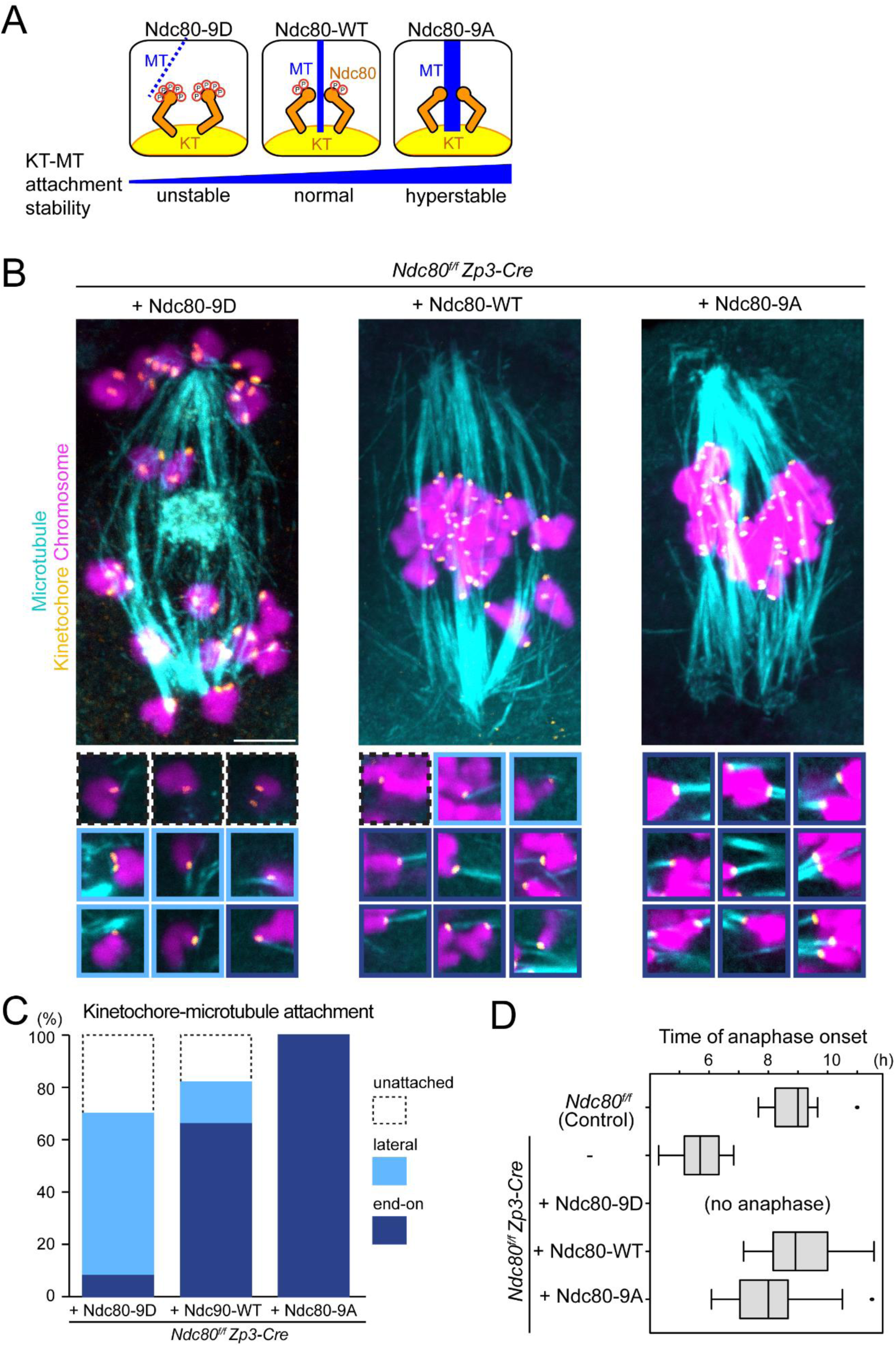
Ndc80 phospho-mutants modify kinetochore–microtubule attachment stability. **(A)** Schematic representation of kinetochore–microtubule attachment. Phosphorylation levels of the Ndc80 complex at the kinetochore are a determinant of the stability of kinetochore (KT)-microtubule (MT) attachments. High phosphorylation levels allow unstable attachments such as lateral attachments. Low phosphorylation levels promote stable attachments such as end-on attachments. Phospho-mimetic and phospho-deficient forms of Ndc80 (S4, S5, T8, S15, S44, T49, S55, S62 and S68 were substituted to aspartic acid “Ndc80-9D” or alanine “Ndc80-9A”, respectively) can be used as tools to manipulate the stability of kinetochore–microtubule attachments. **(B)** Ndc80 phospho-mutants modify kinetochore–microtubule attachment stability. *Ndc80^f/f^ Zp3-Cre* oocytes were collected and microinjected with RNAs of wild-type (Ndc80-WT), phospho-mimetic (Ndc80-9D) or phospho-deficient (Ndc80-9A) forms of Ndc80. Cold-stable microtubules were visualized by immunostaining oocytes 7 hours after NEBD for microtubules (α-tubulin, cyan), kinetochores (ACA, orange), and chromosomes (Hoechst33342, magenta). Z-projection images are shown. Magnified images of kinetochores are categorized: end-on attachments (dark blue box), lateral attachments (light blue box) and unattached (dot-lined box). Scale bar, 10 μm. **(C)** Percentages of kinetochore–microtubule attachments in (B). All kinetochores were analyzed (n=200 kinetochores of 5 oocytes from two independent experiments). **(D)** Anaphase entry timing. *Ndc80^f/f^* (*Ndc80*-intact) oocytes are used as a control. *Ndc80^f/f^ Zp3-Cre* (*Ndc80*-deleted) oocytes expressing Ndc80-WT, Ndc80-9D, or Ndc80-9A were monitored with the chromosome marker H2B-mCherry (n=15, 25, 25, 42 and 55 oocytes, respectively, from at least 3 independent experiments). Time after NEBD. Note that *Ndc80^f/f^ Zp3-Cre* oocytes expressing no Ndc80 construct exhibited an accelerated anaphase onset that is consistent with defects in the spindle checkpoint. Ndc80-9D-expressing oocytes did not undergo anaphase, consistent with spindle checkpoint activation by unstable kinetochore–microtubule attachments. Ndc80-9A-expressing oocytes tended to exhibit an earlier anaphase onset, in agreement with accelerated checkpoint satisfaction by stable kinetochore–microtubule attachments.

**Supplementary Figure 4:**
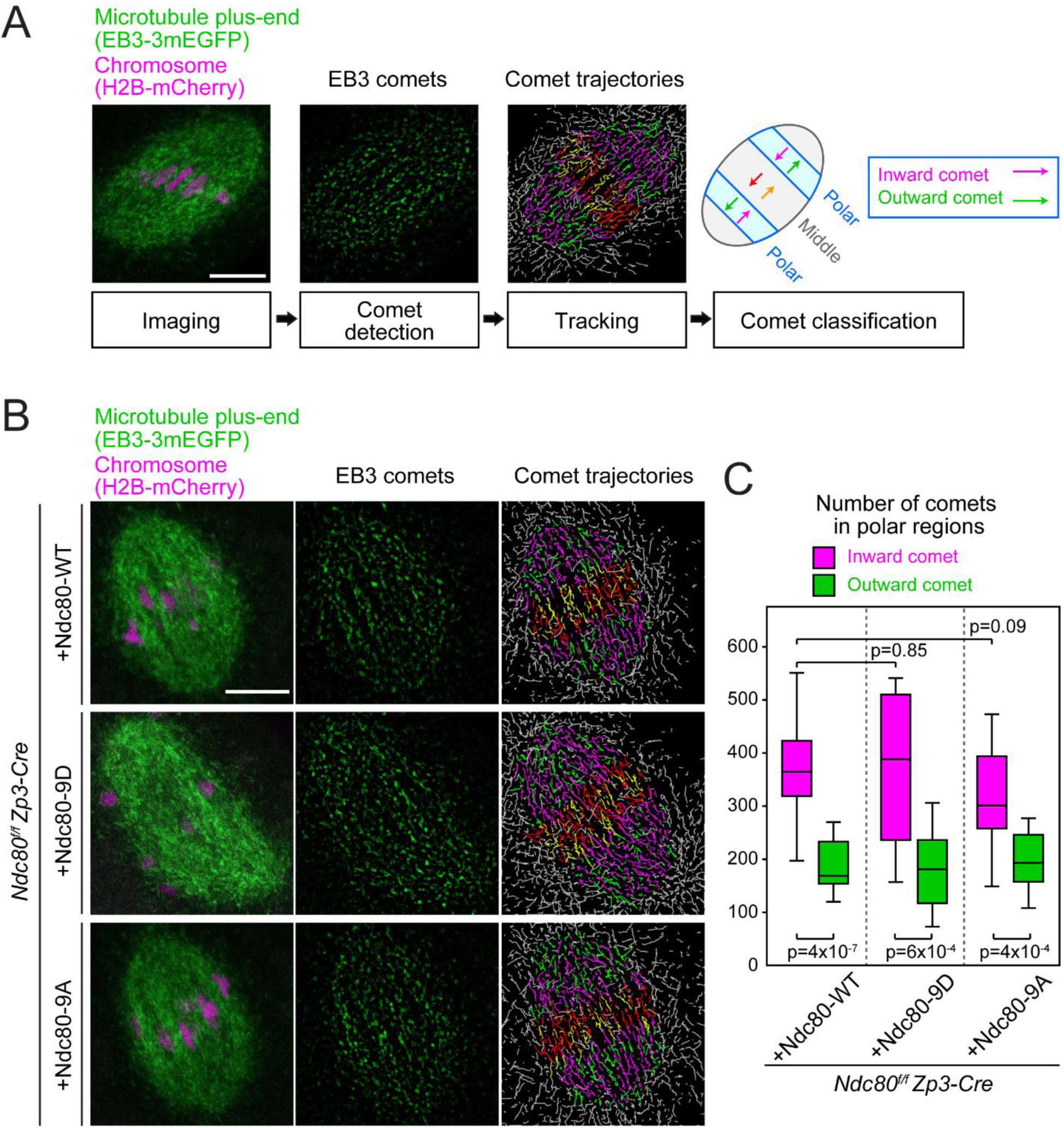
Kinetochore–microtubule attachment stability is not absolutely required for bipolar microtubule organization. **(A)** Experimental pipeline for the quantification of microtubule organization. Oocytes were imaged with the microtubule plus-end marker EB3-3mEGFP (green) and the chromosome marker H2B-mCherry (magenta) at 5 hours after NEBD. The comets of EB3-labeled microtubule plus-ends were detected following image processing for peak enhancement and background subtraction. The comets were tracked to visualize their movements. The spindle was divided into 21 equal regions between the two spindle poles along the spindle axis, and divisions 4–6 and 16–18 were defined as polar regions. In the polar regions, comets moving toward the inside of the spindle were categorized as inward comets (magenta lines), and the others as outward comets (green lines). Inward comets were predominant in the polar regions of the spindle, which represents bipolar microtubule organization. Scale bar, 10 μm. **(B)** EB3 dynamics are largely unaffected by kinetochore–microtubule stability. *Zp3-Cre Ndc80^f/f^* oocytes expressing Ndc80-WT, −9D, or −9A were analyzed as in (A). **(C)** EB3 dynamics are not significantly affected. Total number of comets going inwards and outwards in the polar regions of the spindle (n=14, 13, 16 oocytes from at least 2 independent experiments) is shown. Welch’s t-test was performed. See also Supplementary Movie 8.

**Supplementary Movie 1: MTOC sorting is gradual and continues even in the bipolar-shaped spindle**

Live imaging of MTOC dynamics during meiosis I. Z-projection images of MTOCs (mNG-Cep192, green) and chromosomes (H2B-mCherry, magenta) are shown. Note that central MTOCs are found in the middle of the metaphase spindle. Time in hh:mm. Scale bar, 10 μm. See also Figure 1.

**Supplementary Movie 2: Live imaging of MTOC and kinetochore dynamics**

MTOCs (mNG-Cep192, green) and kinetochores (tdTomato-CENP-C, magenta) in BDF1 oocytes are shown. Time in hh:mm:ss after NEBD. Circles indicate central MTOCs, which were positioned in the middle region of the spindle later than 4 hours after NEBD. Scale bar, 10 μm. See also Figure 2.

**Supplementary Movie 3 and 4: Central MTOCs can be positioned close to kinetochores**

Z-projected images for MTOCs (mNG-Cep192, green) and kinetochores (tdTomato-CENP-C, magenta) are shown. Time in hh:mm:ss after NEBD. Cyan circles follow an MTOC in the middle of the spindle. Magenta circles follow a kinetochore that transiently positions close to the MTOC. Scale bar, 10 μm. See also Supplementary Figure 2.

**Supplementary Movie 5: MTOC dynamics in *Zp3-Cre Ndc80f/f* oocytes expressing Ndc80-WT**

Live imaging of *Zp3-Cre Ndc80^f/f^* oocytes injected with Ndc80-WT. Z-projection images for MTOCs (mNeonGreen-Cep192, green) and chromosomes (H2B-mCherry, magenta) are shown. Time in hh:mm:ss after NEBD. Scale bar, 10 μm. See also Figure 3.

**Supplementary Movie 6: MTOC dynamics in *Zp3-Cre Ndc80f/f* oocytes expressing Ndc80-9D**

Live imaging of *Zp3-Cre Ndc80^f/f^* oocytes injected with Ndc80-9D. Z-projection images for MTOCs (mNeonGreen-Cep192, green) and chromosomes (H2B-mCherry, magenta) are shown. Time in hh:mm:ss after NEBD. Scale bar, 10 μm. See also Figure 3.

**Supplementary Movie 7: Kinetochore–microtubule attachment stability is not required for initiating but is required for terminating spindle elongation**

Live imaging of *Zp3-Cre Ndc80^f/f^* oocytes injected with Ndc80-WT or 9D. Z-projection images show microtubules (EGFP-Map4, green) and chromosomes (H2B-mCherry, magenta). Scale bar, 10 μm. See also Figure 4.

**Supplementary Movie 8: Kinetochore–microtubule attachment stability is not absolutely required for bipolar microtubule organization**

*Zp3-Cre Ndc80^f/f^* oocytes expressing Ndc80-WT, Ndc80-9D, or Ndc80-9A were imaged with the microtubule plus-end marker EB3-3mEGFP (green) and the chromosome marker H2B-mCherry (magenta) at 5 hours after NEBD. The comets of EB3-labeled microtubule plus-ends were detected following image processing for peak enhancement and background subtraction. The comets were tracked to visualize their movements. The spindle was divided into 21 equal regions between the two spindle poles along the spindle axis, and divisions 4–6 and 16–18 were defined as polar regions. In the polar regions, comets moving toward the inside of the spindle were categorized as inward comets (magenta lines), and the others as outward comets (green lines). Inward comets were predominant in the polar regions of the spindle, which represents bipolar microtubule organization. Scale bar, 10 μm. See also Supplementary Figure 4.

## Material and methods

### Ethics Statement

Experiments using mice were approved by the Institutional Animal Care and Use Committee at RIKEN Kobe Branch (IACUC). BDF1 (wild-type), C57BL/6NCrSlc (wild-type), *Ndc80^f/f^* and *Ndc80^f/f^ Zp3-Cre* (C57BL/6 background) (Yoshida et al., in press) female mice, 8–12 weeks of age, were used to obtain oocytes.

### Oocyte culture and in vitro maturation

Mice were injected with 0.1 mL (5 IU) of pregnant mare serum gonadotrophin (PMSG). Oocytes at the GV stage were recovered 44–48 hours later by puncturing ovaries in M2 medium supplemented with 200 nM 3-isobutyl-1-methyl-xanthine (IBMX). Cumulus cells were removed by pipetting the oocytes with a mouth glass pipette. Microinjection was carried out in M2 medium supplemented with IBMX. Resumption of meiotic maturation was induced by washing oocytes with M2 medium at least three times. Oocytes underwent NEBD within 45–75 minutes.

### Microinjection

mRNAs were synthesized using an mMESSAGE mMACHINE T7 kit (Thermo Fisher Scientific) and were purified at a final concentration of 1.0 to 1.5 μg/μL. PCR-based mutagenesis was performed to generate Ndc80-9D and Ndc80-9A, which carry aspartic acid and alanine mutations, respectively, at Ser4, Ser5, Thr8, Ser15, Ser44, Thr49, Ser55, Ser62, and Ser68 of Ndc80. mRNAs were microinjected into GV-stage oocytes in M2+IBMX medium covered with mineral oil, on a heating plate (Tokai-hit) at 37 ºC equipped with an inverted microscope (IX71, Olympus), using micromanipulators (Narishige) and a piezo unit (Prime tech). Microinjection volumes were 4-6 pL for H2B-mCherry (diluted at 1:50 with water), Ndc80 (diluted at 1:4), mNG-Cep192 (diluted at 1:20), tdTomato-CENP-C (diluted at 1:17), and EB3-3mEGFP (diluted at 4:5). Oocytes were incubated for 3 hours in M2+IBMX medium before resumption of meiotic maturation was induced.

### Live imaging of MTOCs, spindles and kinetochores

Oocytes were imaged using a Zeiss LSM780 confocal microscope equipped with a 40x water-immersion C-Apochromat 1.2 NA objective (Zeiss) at 37ºC in M2 medium. The microscope was controlled using Zen black software (Zeiss). Oocytes were tracked using the macro provided by Dr. Jan Ellenberg at EMBL Heidelberg (Politi et al., 2018). Pinhole size was adjusted to acquire 1.5-µm-thick confocal sections. Z-stacks were acquired with 0.9–2.5 µm intervals between z-slices. mEGFP and mNeonGreen were excited with a 488-nm argon laser. mCherry and tdTomato were excited using a 561-nm DPSS laser.

### Immunostaining

Oocytes were fixed with 1.6% paraformaldehyde (pre-warmed at 37ºC) for 30 minutes at room temperature, followed by permeabilization for 15 minutes in 0.2% PBS plus 0.1% Triton-X100 (PBT). In experiments to visualize cold-stable microtubules, we pre-cooled 200 μL M2 medium in a 1.5 mL tube on ice for at least 20 minutes, transferred oocytes into the medium and then incubated them for 7 minutes on ice before fixation. Oocytes were then blocked with 3% BSA in 0.1% PBT at room temperature for at least 15 minutes (up to 2 hours) and incubated them at 4ºC overnight with primary antibodies. Oocytes were subsequently incubated with secondary antibodies for 2 hours at room temperature. Hoechst33342 (Molecular Probes) was added at 1:2000 in PBS + 1% BSA. The primary antibodies used were anti-pericentrin (mouse, 611814, BD Transduction Laboratories, 1:500), anti-alpha-Tubulin (rat, YL1/2, MCA77G, Bio-Rad, 1:2000 (Figure 5) or 1:10000 (other figures); or mouse, DM1A, T6199, Sigma, 1:500) and ACA (human, 15-234, Antibodies Incorporated, 1:200 (Figure 5) or 1:500 (other figures). Secondary antibodies used were Alexa Fluor 488 goat anti-mouse IgG (H+L) (A11029); goat anti-rat IgG (H+L) (A11006); Alexa Fluor 555 goat anti-mouse IgG (H+L) (A28180); goat anti-human IgG (H+L) (A21433); Alexa Fluor 647 donkey anti-mouse IgG (H+L) (A31571) (Molecular Probes, 1:500 (Figure 5) or 1:200 (other figures)).

Fixed oocytes were imaged using a Zeiss LSM780 confocal microscope equipped with a 40x water-immersion C-Apochromat 1.2 NA objective (Zeiss) at a constant temperature (25ºC). The microscope was controlled using Zen Black software (Zeiss). Pinhole size was adjusted to acquire 1-µm-thick optical sections. Z-stacks were acquired with 0.5 µm intervals between z-slices. A 405-nm diode laser, a 488-nm argon laser, a 561-nm DPSS laser, a 633-nm HeNe laser were used for exciting Alexa Fluor 488, 555 and 633/647, respectively.

### Image analysis

MTOC, spindle and chromosome signals were reconstructed into 3D using Imaris software (Bitplane). For every time point, the spindle axis was manually defined based on the spindle shape or MTOC distribution. For earlier time points before spindle axis establishment, the same axis as that of early metaphase (immediately after spindle axis establishment) was used. To generate kymographs and density maps of MTOCs and chromosomes, the intensities of their signals or the volumes of 3D-reconstructed objects were projected onto the spindle axis for each time point, using Fiji (Schindelin et al., 2012). In kymographs, the positions of two poles were manually determined, and were used to define the center of the spindle axis. In the density maps, the center of mass of chromosomes was used as the center of the spindle.

### MTOC and kinetochore tracking pipeline

MTOC and kinetochore positions were detected using the spot function of Imaris software (Bitplane) and were manually verified. These positions were then used for the tracking function of Imaris, and the resultant tracks were manually verified. The spindle axis for each time point was defined based on the distribution of MTOCs. The spot positions of the tracks and the orientation of the spindle axis were exported to R software. The positions of MTOCs and kinetochores along the spindle axis were calculated with R. The distance between each kinetochore pair and MTOCs was calculated in 3D with R. Individual MTOC and kinetochore pairs trajectory could then be extracted using R for each middle MTOC. For experiments to determine the origin of central MTOCs, the trajectories were manually verified with 3D-reconstructed images in Imaris.

### EB3 imaging and tracking pipeline

Oocytes microinjected with mRNAs for 3mEGFP-EB3 and H2B-mCherry were imaged on an LSM780 microscope, equipped with a 40x C-apochromat water objective (1.2 NA) at 37ºC at 5 hours after NEBD. Single frames were acquired consecutively without waiting time. Scan time was 782 msec for one frame (512×512 pixels). For peak enhancement and background subtraction, we subtracted the image blurred with a strong Gaussian filter (sigma 8) from the same original image blurred with a light Gaussian filter (sigma 1.2). Tracking of EB3 comets was done automatically using the Trackmate plugin (Tinevez et al., 2017) in Fiji. To determine the position and direction of EB3 comets relative to the spindle, we manually fit an ellipse on the spindle using ImageJ, which was used to determine the position and orientation of the spindle axis and equator. The data for EB3 tracks and the spindle were then analyzed using R.

## Acknowledgments

We thank M. Schuh for providing a Cep192 construct, J. Ellenberg for providing a macro for automated microscopy, the imaging and genome analysis and animal facilities of RIKEN Kobe for technical support. We also thank our laboratory members. A.C. was partly supported by JSPS Postdoctoral Fellowship. This work was supported by the research grants MEXT/JSPS KAKENHI JP16H06161/JP16H01226/JP18H05549 to T.S.K.; JSPS KAKENHI JP17K15069/JP19K06682 to S.Y.; and by RIKEN BDR.

## Author Contributions

A.C. designed and performed almost all experiments, analyzed and interpreted the data, and wrote the manuscript. S.Y. performed experiments shown in Figure 1E, 4, 5, and Supplementary Figure 1. T.S.K. designed, conceptualized, and supervised the project, interpreted the data, and wrote the manuscript.

